# Spiking Neuron-Astrocyte Networks for Image Recognition

**DOI:** 10.1101/2024.01.10.574963

**Authors:** Jhunlyn Lorenzo, Juan-Antonio Rico-Gallego, Stéphane Binczak, Sabir Jacquir

## Abstract

From biological and artificial network perspectives, researchers have started acknowledging astrocytes as computational units mediating neural processes. Here, we propose a novel biologically-inspired neuron-astrocyte network model for image recognition, one of the first attempts at implementing astrocytes in Spiking Neuron Networks (SNNs) using a standard dataset. The architecture for image recognition has three primary units: the pre-processing unit for converting the image pixels into spiking patterns, the neuron-astrocyte network forming bipartite (neural connections) and tripartite synapses (neural and astrocytic connections), and the classifier unit. In the astrocyte-mediated SNNs, an astrocyte integrates neural signals following the simplified Postnov model. It then modulates the Integrate-and-Fire (IF) neurons via gliotransmission, thereby strengthening the synaptic connections of the neurons within the astrocytic territory. We develop an architecture derived from a baseline SNN model for unsupervised digit classification. The Spiking Neuron-Astrocyte Networks (SNANs) display better network performance with an optimal variance-bias trade-off than SNN alone. We demonstrate that astrocytes promote faster learning, support memory formation and recognition, and provide a simplified network architecture. Our proposed SNAN can serve as a benchmark for future researchers on astrocyte implementation in artificial networks, particularly in neuromorphic systems, for its simplified design.

## 1 Introduction

Twenty years ago, Araque, et al. introduced the concept of the tripartite synapse, where the astrocyte acts as the third component in synaptic communication [1]. An astrocyte, through its peripheral processes, enwraps the synaptic space between the preand postsynaptic neurons, as shown in Figure 1. The presynaptic neuron encodes information in the action potential (AP) pattern (in the spiking activity of its membrane potential), and sends it to the neighboring neurons through the neurotransmission process, wherein glutamate molecules (Glu^−^) are released into the synaptic space [2]. A fraction of synaptic Glu^−^ activates the postsynaptic Glu^−^ receptors (GluRs), or diffuses to the extrasynaptic space and activates the metabotropic GluRs (mGluRs) on the astrocytic processes [3]. Therefore, the presynaptic neuron also modulates the activity of the astrocyte by indirectly producing inositol triphosphate (IP_3_) molecules that results in the increase of calcium concentration ([Ca^2+^]) in the intracellular space [4]. When the intracellular [Ca^2+^] increases above a threshold (Ca^2+^ spike), the astrocyte releases extrasynaptic Glu^−^. In this case, the astrocyte modulates the postsynaptic neuronal components by activating extrasynaptic GluRs [5]. The postsynaptic neuron integrates the fast and slow excitatory signals produced by neuron- and gliotransmission, respectively. Therefore, the firing patterns of the presynaptic neuron and the astrocytic process work in conjunction to regulate postsynaptic signals. The tripartite synapse concept implies that astrocytes not only provide structural support to neurons, but also serve as pivotal elements in neuronal communication [6].

**Fig. 1:**
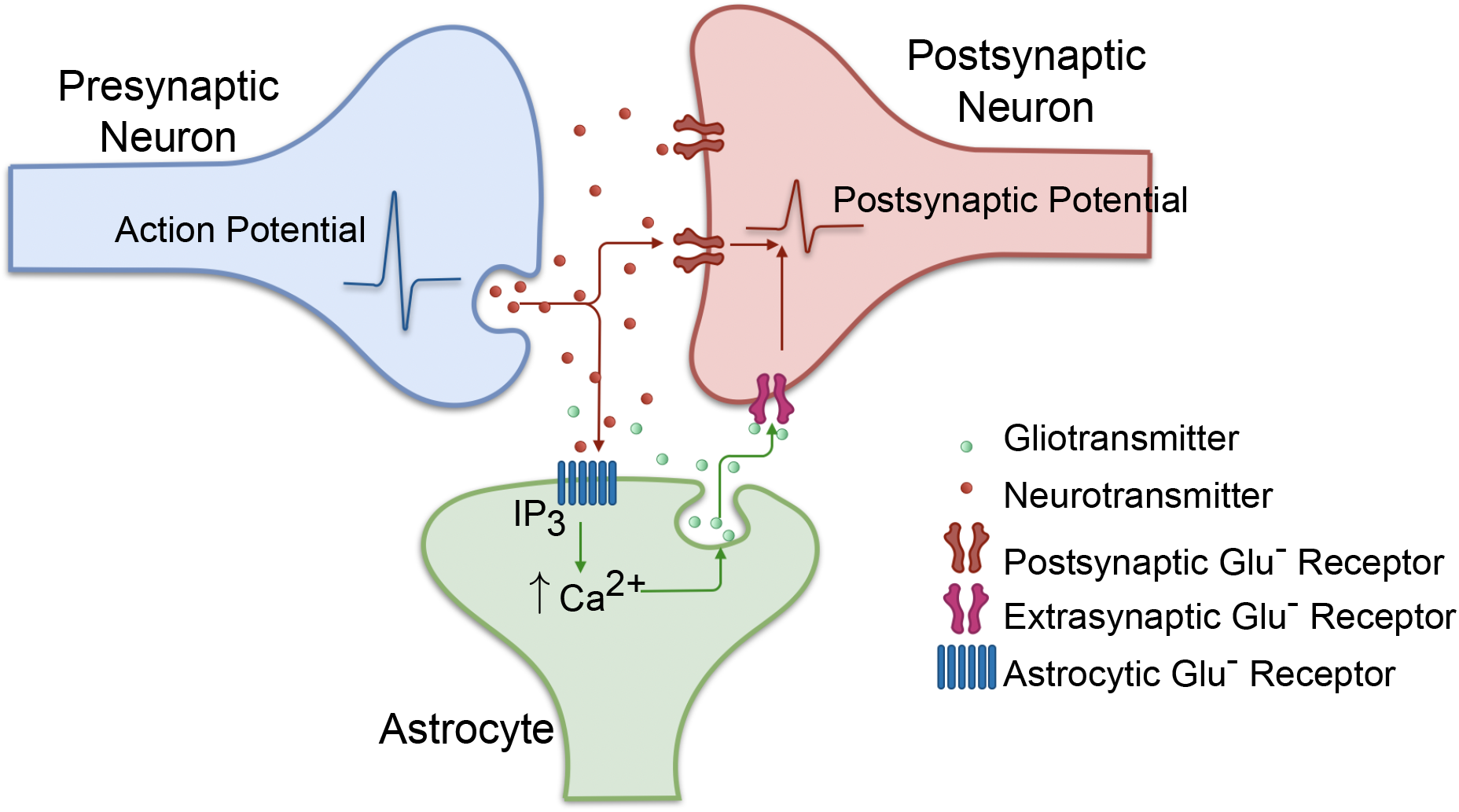
The tripartite synapse.

Extending the scope of investigation beyond the tripartite synapse to encompass the entirety of the astrocytic cell reveals that a single astrocyte can enwrap numerous synapses originating from multiple pre- and postsynaptic neurons, as illustrated in Figure 2 [7]. Ca^2+^ signalling, which is primarily triggered by IP_3_ production, is the main signalling mechanism of astrocytes [8]. The astrocytic processes can be viewed as distinct compartments, each having independent Ca^2+^ signaling mechanisms. These compartments can communicate with one another via the intracellular diffusion of Ca^2+^ and IP_3_ into the soma or adjacent compartments, and vice versa [9, 10]. Given the complex architecture of astrocytes, there exist different pathways by which astrocytes can modulate neighboring neurons: (1) A tripartite synapse that remains mostly inactive due to the absence of presynaptic activity can be activated by Ca^2+^ that diffuses from both the astrocytic soma and other active compartments [11]. (2) All tripartite synapses can be activated synchronously during a global Ca^2+^ elevation caused by the integration of Ca^2+^ from all compartments [12].

**Fig. 2:**
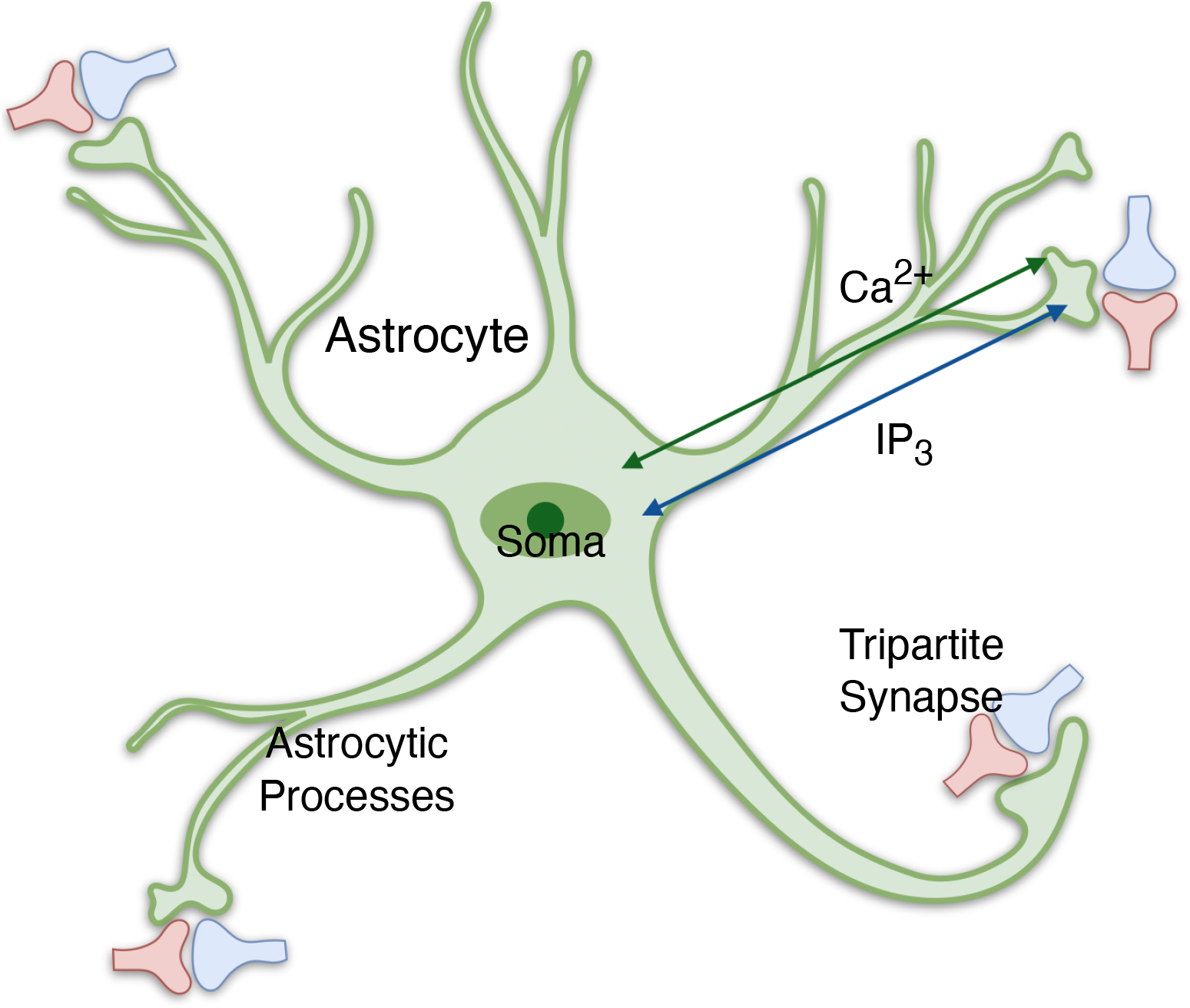
An astrocyte forming tripartite synapses

Astrocytes are key players in synapse formation [13] and in enhancing neuronal synchrony [14] by regulating stochastic neuronal spiking. Another advantage of astrocytes in the neural network is their function in memory storage−in maintaining long-term hippocampal potentiation [15]. However, the use of astrocytes as components in artificial neural networks is a more recent concept. Why should astrocytes be considered as computational units in neural networks? Analogous to the McCulloch-Pitts neuron model [16], which was inspired by biological neurons and laid the foundation for modern artificial neural networks, astrocytes demonstrate several similarities to neurons, despite their relatively slower signaling dynamics. Both neurons and astrocytes can modulate synaptic weights with neurons in adjacent layers via neurotransmission and gliotransmission [10]. Neurons integrate synaptic inputs and generate spikes upon reaching a specific threshold, whereas astrocytes integrate Ca^2+^ signals and spike once the Ca^2+^ threshold has been attained [3]. In addition, astrocytes also form astrocytic networks in which each unit communicates via Ca^2+^ wave propagation [17].

The third generation of neural networks, known as Spiking Neural Networks (SNNs), is becoming increasingly competitive with their Artificial Neural Network (ANN) counterparts. This is primarily due to their ability to emulate brain-like dynamics and perform event-based computations [18–20]. Inspired by biological neural mechanisms, SNNs employ spike-based signaling to perform brain-like computations, integrate and represent spatiotemporal information, process sparse and asynchronous signals, and execute parallel processing [21]. Similar to actual neural circuits, SNNs offer low-power consumption, analog computation, rapid inference, online learning, and event-driven processing. The temporal dimensions of SNNs, in particular, make them prime candidates for real-world applications and neuromorphic hardware implementations compared to other deep neural networks [22].

While models of SNNs closely mimic brain processes, there are some drawbacks. Training using benchmarks such as MNIST (Modified National Institute of Standards and Technology) [23] and ImageNet [24] indeed yields a lower accuracy than ANNs, which can be attributed to converting the frame-based images into rate-coded information. Besides, there is a lack of training algorithms for spiking networks. Training SNNs means dealing with the asynchronous and discontinuous nature of spikes, making the application of current differentiable backpropagation techniques quite challenging [18, 22]. Here, we draw inspiration from biological processes to address these issues, particularly from the astrocyte-mediated synapse−the tripartite synapse. This study is a proof of concept that astrocytes can be employed in image recognition to improve SNNs performance. While recent examples include the bio-inspired neuron-astrocyte network for image classification [25, 26] and the unsupervised network with self-repair in the event of synaptic failure [27], astrocytes employed here are for neural support rather than as computational units.

This study presents a novel approach for improving image recognition performance through the integration of astrocytes in spiking neural networks. The proposed Spiking Neural Network with Astrocyte Network (SNAN) architecture includes simplified neuron and astrocyte models, allowing for efficient and accurate information processing, and employs an unsupervised learning scheme based on spike-timing-dependent plasticity (STDP). Subsequently, the network was trained using the MNIST dataset to determine the optimal hyperparameters concerning network architecture and learning. We optimized network parameters such as synaptic coupling strengths and firing thresholds to enhance network performance. Upon training the network, we evaluate its accuracy in predicting the output digits. Finally, we assess the influence of astrocytes on network performance. The results show that astrocytes significantly improve network performance and facilitate faster learning compared to conventional spiking neural networks (SNNs) alone.

## 2 SNAN Architecture

This section describes the tripartite synaptic connection, the neuron and astrocyte layers, and the corresponding network architecture. It also includes the learning algorithm used to train the network, the simulation process and the performance analysis method.

### 2.1 Tripartite Synapse Model

In the proposed network architecture, as depicted in Figure 3, the tripartite synapses comprise Poisson spiking presynaptic neurons (*n*^1^) that are coupled to the astrocyte (*a*^1^) and the postsynaptic neuron (*n*^2^) via the synapses (*s*^1^). The spiking rate of the Poisson neurons depends on the input image pixel intensity, which ranges from 0 for black to 31.875 Hz for white pixels. The rate is chosen to ensure that the presynaptic inputs do not oversaturate the astrocytic dynamics, while generating a postsynaptic spike. The postsynaptic neuron integrates three types of inputs: (1) fast excitatory inputs arising from neurotransmission, (2) slow excitatory inputs due to gliotransmission, and (3) inhibitory inputs originating from the interneurons (*m*^2^). The postsynaptic neuron potential (*v*) in mV is governed by the adaptive integrate-and-fire (AIF) neuron model [28] given as follows:

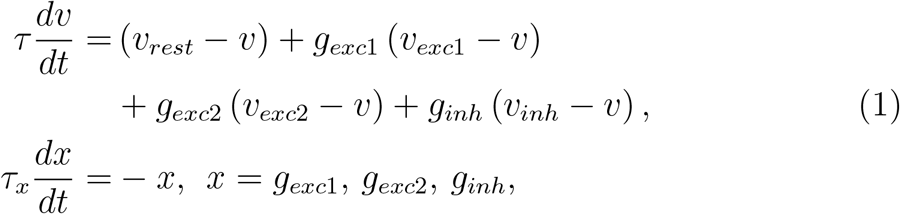

The conductance (in nS) of the fast excitatory, slow excitatory, and inhibitory synapses are equal to the weight summations *g*_*exc*1_, *g*_*exc*2_, and *g*_*inh*_, respectively, and where *x* defines their conductance decay through time *t*. The parameter *v*_*rest*_ is the membrane resting potential while *v*_*exc*1_, *v*_*exc*2_, and *v*_*inh*_ are the reversal potentials of the excitatory and inhibitory synapses. The constant parameters, *τ* and *τ*_*x*_, are the time constants of the membrane potential and the synapses, respectively. When the AIF neuron membrane potential *v* crosses its spiking threshold *θ*, it generates a spike, then immediately resets to a level equal to *v*_*reset*_, and remains here until the refractory period, *τ*_*ref*_, ends. During training, the adaptive threshold increases by *α*^*′*^ after a spike and decays following the exponential model

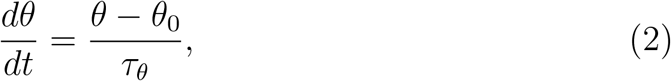

where *θ* = *v*− *v*_*reset*_ and *τ*_*θ*_ is the time constant.

**Fig. 3:**
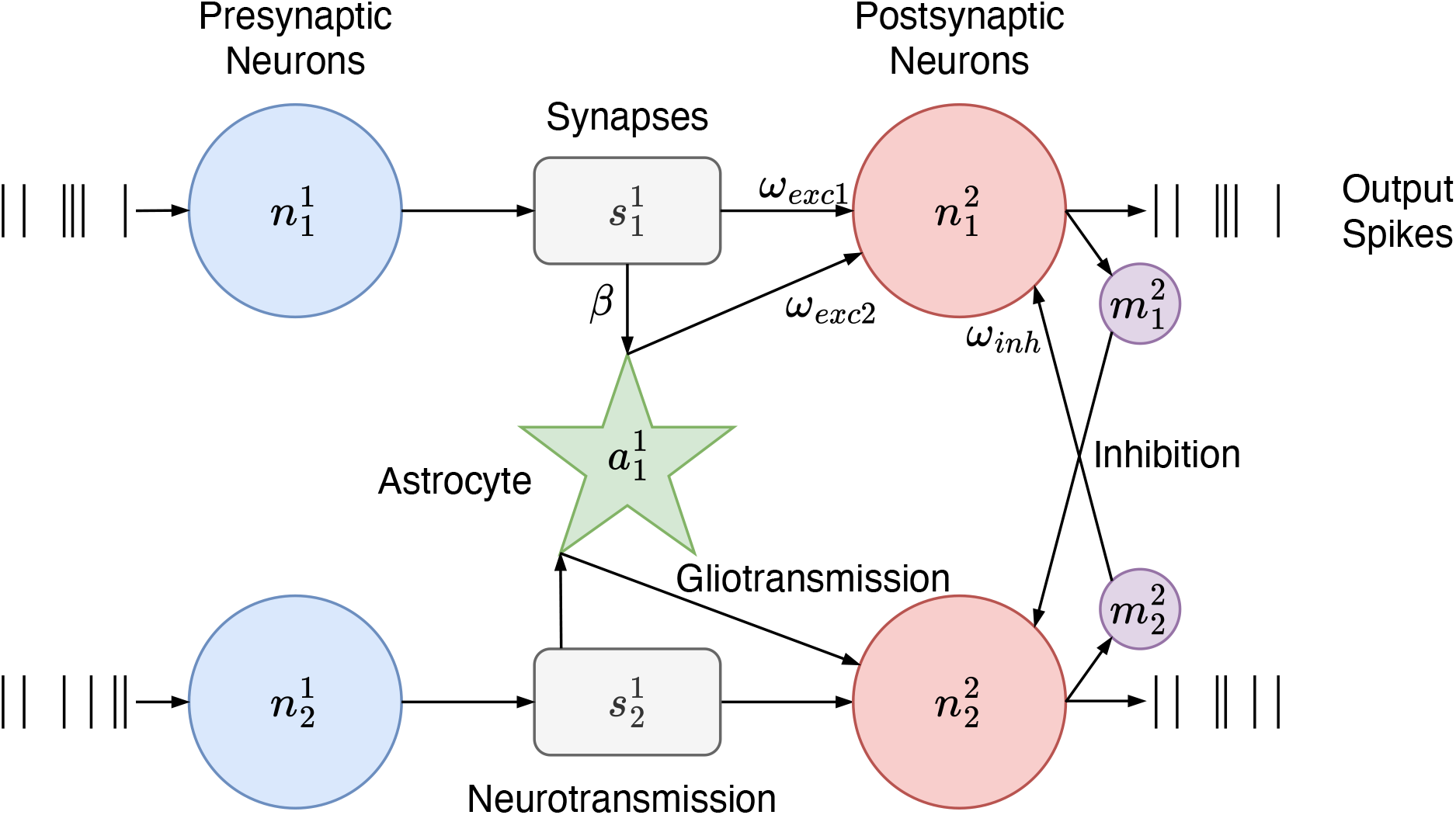
The tripartite synapse connections.

The postsynaptic neuron layer consists of lateral inhibition for maintaining homeostasis and signaling the postsynaptic neuron activity to its neighbors. Here, the postsynaptic neuron signals its spiking activity to the IN, and in return, the IN then inhibits the activity of the other neurons, generating competition among neurons. The interneurons follow the IF model in Equation 1 but without the adaptive property *θ* and the *g*_*exc*2_ and *g*_*inh*_ conductances, while *g*_*exc*1_ changes with respect to the postsynaptic neuron spiking activity. Refer to Table 1 for the list of parameters of the postsynaptic excitatory and inhibitory neurons.

**Tab. 1:** List of parameters of the postsynaptic neurons with lateral inhibition.

| Parameter | Value | Description |
| --- | --- | --- |
| *Postsynaptic Excitatory Neurons* |  |  |
| $v_{rest}$ | -65 mV | Membrane resting potential |
| $v_{exc1}$ | 0 mV | Fast synaptic input reversal potential |
| $v_{exc2}$ | 0 mV | Slow synaptic input reversal potential [29] |
| $v_{inh}$ | -100 mV | Inhibitory synaptic reversal potential |
| $\tau$ | 100 ms | Membrane time constant |
| $\tau_{exc1}$ | 1 ms | Excitatory synapse (fast) time constant |
| $\tau_{exc2}$ | 600 ms | Excitatory synapse (slow) time constant [29] |
| $\tau_{inh}$ | 2 ms | Inhibitory synapse time constant |
| $v_{reset}$ | -65 mV | Reset potential |
| $v_{thresh}$ | -52 mV | Spiking threshold level |
| $t_{ref}$ | 5 ms | Refractory period |
| $\theta_0$ | 20 mV | Spiking threshold offset |
| $\tau_\theta$ | 107 ms | Spiking threshold time constant |
| $\alpha'$ | 0.05 mV | Increase in spiking threshold |
| *Postsynaptic Inhibitory Neurons* |  |  |
| $v_{rest}$ | -60 mV | Membrane resting potential |
| $v_{exc1}$ | 0 mV | Synapse reversal potential |
| $\tau$ | 100 ms | Membrane time constant |
| $\tau_{exc1}$ | 1 ms | Excitatory synapse time constant |
| $v_{reset}$ | -45 mV | Reset potential |
| $v_{thresh}$ | -40 mV | Spiking threshold |
| $t_{ref}$ | 2 ms | Refractory period |

Unless otherwise stated, the parameters are taken from Diehl and Cook [30].

The astrocyte function as a point process integrating the presynaptic inputs, following the simplified astrocytic model proposed by Postnov et al. [31, 32]. The synaptic coupling variable *z* spikes to 1, synchronous with the presynaptic neuron spiking, and then immediately inactivates by decaying to 0, following the simplified model

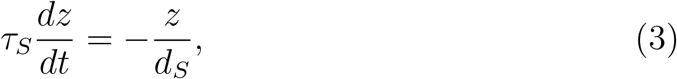

where *d*_*S*_ controls the relaxation of *z* and *τ*_*S*_ is the synaptic delay. The variable *z* triggers the astrocytic IP_3_ production, *S*_*m*_, described as

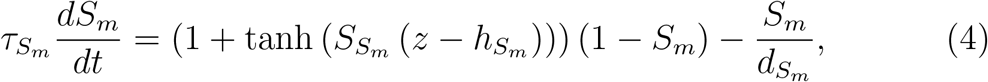

where the parameters 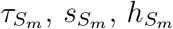. and 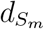 control the time scale, steepness of activation of the sigmoid function, threshold values, and the deactivation rate, respectively. Then, the increase in *S*_*m*_ results in Ca^2+^ elevation, *c*, described by the system shown in Equation 5.

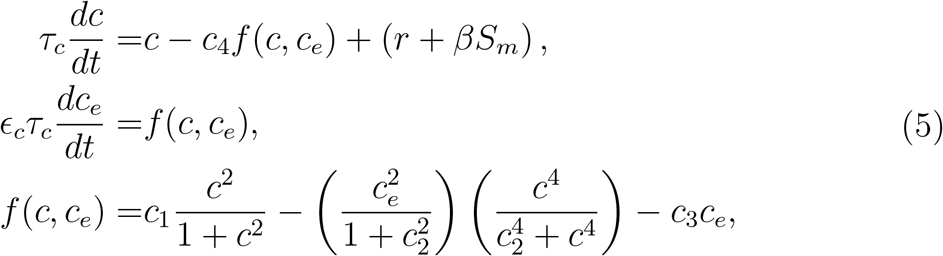

where *c*_2_ denotes the Ca^2+^ concentration in the ER, the constant parameters *ϵ*_*c*_, *τ*)_*c*_ together define the characteristic time for Ca^2+^ oscillations. The factor *β* controls the variable *S*_*m*_ and *r* controls the initial state of Ca^2+^ oscillation. The nonlinear function *f* (*c, c*_*e*_) describes the Ca^2+^ exchange between the cytoplasm and the ER.

The astrocyte then integrates the presynaptic signals from multiple synaptic connections, where *β* is the control parameter defining the magnitude of influence of the presynaptic neuron activity on the astrocytic Ca^2+^. The modified Ca^2+^ model [32] is defined as

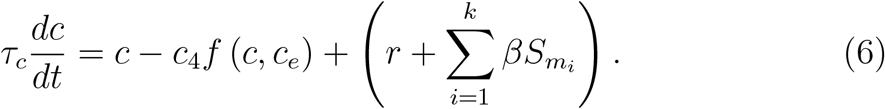

When the astrocytic [*Ca*^2+^], *c*, increases beyond the threshold, *c*_*thresh*_, it triggers the release of extrasynaptic Glu^−^, activating slow excitatory synapses of the coupled postsynaptic neurons. After the Glu^−^ spike, the astrocyte stays in the refractory period provided that *c* stays above the threshold. The astrocytic parameters are listed in Table 2.

**Tab. 2:** Simplified astrocytic model parameters.

| Parameter | Value | Description |
| --- | --- | --- |
| *Neuron-astrocyte coupling* |  |  |
| $\tau_S$ | 10 ms | Synaptic delay time constant |
| $s_S$ | 1 | Steepness of activation |
| $h_S$ | 0 | Activation control |
| $d_S$ | 3 | Relaxation control |
| *IP<sub>3</sub> production* |  |  |
| $\tau_{Sm}$ | 100 ms | Time constant |
| $s_{Sm}$ | 100 | Steepness of activation |
| $h_{Sm}$ | 0.45 | Threshold |
| $d_{Sm}$ | 3 | Deactivation rate |
| *Astrocytic Ca<sup>2+</sup>* |  |  |
| $\epsilon_c$ | 0.04 | Characteristic time control |
| $c_1$ | 0.13 | Ca<sup>2+</sup> oscillation control parameters |
| $c_2$ | 0.9 | |
| $c_3$ | 0.004 | |
| $c_4$ | $2/\epsilon_c$ | |
| $r$ | 0.31 | Ca<sup>2+</sup> oscillation initial state |
| $\tau_c$ | 600 ms | Time constant for Ca<sup>2+</sup> oscillation |

The parameters are taken from Postnov et al. [31, 32].

### 2.2 Network Architecture

The digit recognition process follows the network architecture presented in Figure 4. The process starts from the image preprocessing unit, where the input or sample image is converted into rate-coded signals, followed by unsupervised learning by the SNAN, and lastly, based on the SNAN output spike patterns, the machine learning (ML) model predicts the output digit. Given a sample digit from the MNIST dataset, a 28*×*28-pixel image of a handwritten digit is converted into a 784*×*1 vector of pixel intensities. One pixel intensity corresponds to the spiking frequency that drives a Poisson neuron in the input layer of the SNAN.

**Fig. 4:**
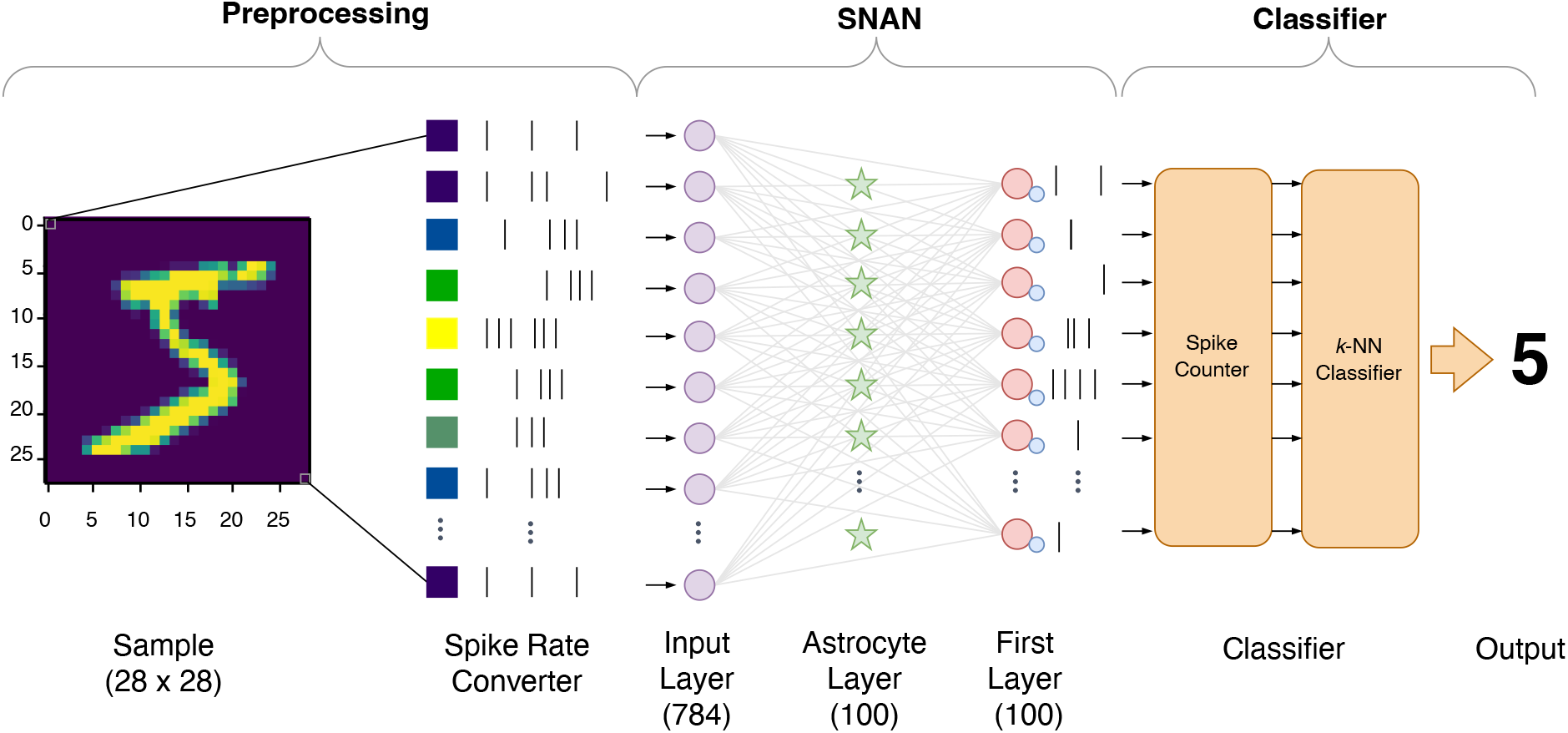
Network architecture for image recognition is a three-stage process comprising an input preprocessing unit, a SNAN learning unit, and a classifier unit.

In the spiking network, 784 Input layer neurons form 78 400 fully connected (dense) synapses with the neurons in the First Layer. There is a 1:1 ratio between the number of excitatory and inhibitory neurons in the First Layer, where the forward connection (from the excitatory to the inhibitory) is a one-to-one topology. At the same time, the lateral inhibitions form 9900 inhibitory synapses, where an IN connects with all excitatory neurons in the same layer except with the one in the forward connection. The astrocytic layer between the two neuronal layers, also following a 1:1 ratio with the First layer neurons, forms the tripartite synapses by connecting with the synapses rather than directly with the input layer neurons. Because an astrocyte is coupled with the synapses, it can receive multiple presynaptic inputs from the same Input layer neuron and increase its influence on a First layer neuron by modulating multiple tripartite synapses coupled to that postsynaptic neuron.

In the classifier unit, the spike counter converts the output spiking patterns of the First layer excitatory neurons into a vector of whole numbers corresponding to the number of spikes in each neuron. This vector gives the set of input features for the classifier. Our simulation analysis suggests that the cosine *k*-NN classifier using 5-fold cross-validation yields a faster and more accurate classification performance than the other available machine learning classifier. Then lastly, the network predicts the output from ten classes labeled from 0 to 9. This paradigm requires that the SNAN generates a stable spiking pattern for each input class for the network to recognize and differentiate each input pattern.

### 2.3 Spike-Timing-Dependent Plasticity

A form of long-term plasticity called spike-timing-dependent plasticity (STDP) addressed the temporal issue in Hebbian plasticity [33–35]. Rather than correlated inputs driving the synaptic plasticity, the spiking activities of neurons within a temporal window define the direction of synaptic plasticity, and the relative timing between the pre- and postsynaptic firing determines the change in the synaptic weights. Therefore, the SDTP learning rule is suitable for training the SNAN due to the significant difference between the neuronal and astrocytic time scales.

In the STDP rule following Diehl and Cook model [30], the increase in synaptic weight includes a weight-dependent term so that during postsynaptic activation, the weight increases only after the presynaptic activation. Let the traces *α*_pre_, *α*_*post*1_, and *α*_*post2*_ decay exponentially by

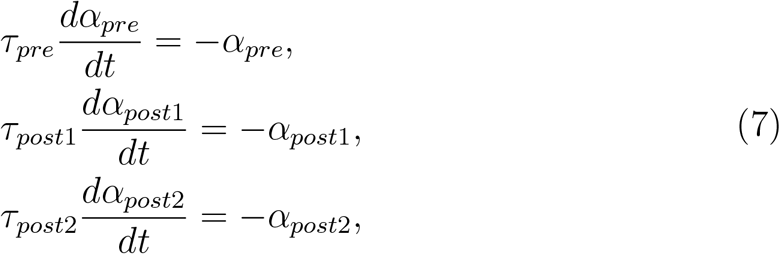

where *τ*_*pre*_, *τ*_*post1*_, and *τ*_*post*2_ are the decay constants equal to 20 ms, 20 ms, and 40 ms, respectively. On presynaptic activation, the weights are updated following the rule

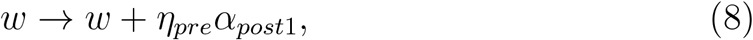

while on the postsynaptic activation,

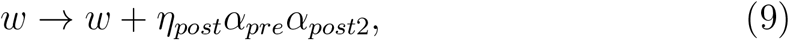

where *η*_*pre*_ and *η*_*post*_ are the pre- and postsynaptic rates, respectively. The excitatory-to-excitatory connections use the pre- and postsynaptic spiking weight update rules, while astrocyte-postsynaptic coupling only increases postsynaptic weights during astrocytic activation. Due to the significant time scale difference between neurons and astrocytes, the slow synapse conductance depletes to zero when the postsynaptic neuron fires continuously. Moreover, the excitatory-to-inhibitory and the inhibitory-to-excitatory Δ*w* are constants equal to 17 and 10.4, respectively.

### 2.4 Simulation Method and Performance Analysis

Here, we used the Brian2 simulator [36], in conjunction with the python programming language, as it explicitly describes models in a high-level form by writing the differential equations directly in the code, and allows us to efficiently model and to integrate astrocytes into the network.

The MNIST dataset [23] contains 60 000 training, 5000 validation, and 5000 test images. Each image is fed into the proposed network one at a time for 350 ms, followed by a resting time of 150 ms, to ensure that the network activity due to the previous image will not overlap with the new sample; therefore, one sample takes 500 ms to pass through the SNAN. The network continuously updates its weights via the STDP learning rule and the neuronal firing thresholds during training. We divide one epoch (containing all the training images) into batches of 1000 samples each and then normalize the weights after each batch. After each epoch, the validation set is fed into the network using the learned parameters.

We simulated different spiking network configurations to determine the hyperparameters that lead to optimum network performance: one SNN and three SNANs (SNAN,1, SNAN2, SNAN3). All networks have the same number of neurons and astrocytes (except in SNN), while the number of synaptic connections with astrocytic coupling increases from 10% to 60% (increment of 10%) of the total synapses, where each astrocyte receives the same number of presynaptic inputs. The set of hyperparameters is listed in Table 3. The simulations were performed using the high-performance computing resources from the Barcelona Computing Center and the University of Burgundy Computing Center.

**Tab. 3:** Set of hyperparameters per configuration.

| Hyperparameters | SNN | SNAN1 | SNAN2 | SNAN3 |
| --- | --- | --- | --- | --- |
| $\eta_{exc1_{pre}}$ | 0.0001 | 0.0001 | 0.0001 | 0.0001 |
| $\eta_{exc1_{post}}$ | 0.01 | 0.01 | 0.01 | 0.01 |
| $\eta_{exc2_{pre}}$ | - | 0.000001 | 0.00001 | 0.000001 |
| $\eta_{exc2_{post}}$ | - | - | - | - |
| $w_{exc1_{max}}$ | 1 | 1 | 1 | 1 |
| $w_{exc2_{max}}$ | - | 0.001 | 0.01 | 0.001 |
| $\beta$ | - | 0.00005 | 0.00005 | 0.0001 |

### 2.5 Performance analysis

The classifier accuracy defines how well the system groups and distinguishes the spiking patterns generated by the SNAN from one class to another. Therefore, the classification performance depends on the extent to which the SNAN learned the rate-coded images. During training, we determine the accuracy after each epoch. Then, using the learned parameters after each epoch, we feed the validation set into the network for prediction and compare the results with the training accuracy. Therefore, the parameters chosen for the test network simulations are the learned parameters of the epoch giving the maximum average accuracy between the train and validation sets.

## 3 Results

We simulated the spiking networks and set the SNN as the baseline of the SNAN activities. Here, we focus on the influence of astrocytes in network activities. The minimum numbers of neurons for conventional MNIST classification networks are 784 and 100 in the Input and First layers, respectively. Here, the number of astrocytes in all networks is equal to the number of First layer excitatory neurons (1:1 ratio); however, the number of synapses covered by astrocytes increases by 10% of the total number of Input to First layer synaptic connections. A higher number of neurons results in more synapses. Moreover, astrocytes create two more types of synaptic coupling (presynaptic neuron-to-astrocyte and astrocyte-to-postsynaptic neuron shown in Figure 3), resulting in a more complex network that requires prolonged simulations.

The SNAN design is a highly iterative process, requiring numerous trials before arriving at the possible set of hyperparameters shown in Table 3. The neuronal and neuron-to-neuron synaptic hyperparameters are consistent in all the networks. Based on our initial simulations, these parameters lead to regular neuronal spiking. The Poisson neuron changes its spiking rate every 500 ms. We also determined that the Poisson neuronal firing rate lies between 0 to 32 Hz (proportional to the pixel intensity from 0 to 255) to prevent oversaturation of astrocytic *S*_*m*_ (IP_3_ pathway) and avoid overexcitation of astrocytic Ca^2+^ dynamics, *c*.

### 3.1 Tripartite Synapse Dynamics

Figure 5 is an overview of the activities of a single tripartite synapse without STDP and wherein the astrocyte only receives a presynaptic input and modulates one postsynaptic neuron. The presynaptic neuron generates 5 Hz and regularly spaced spikes. The synaptic coupling variable *z* increases to 1 simultaneously with the input neuron spikes and then exponentially decays back to 0, replicating the neurotransmitter release and recycling process. In addition, the IP_3_ pathway (*S*_*m*_) also increases and decays with *z*, and whose peak amplitudes stabilize at 0.54. The Ca^2+^ variable *c* generates spikes, upon crossing the 0.4 threshold, with a peak amplitude equal to 1.29 with a pulse duration of 58 ms. After activation, the astrocyte is in the refractory period provided that *c >* 0.4. A single fast excitatory synaptic input generates a postsynaptic spike, *v*, with −64.38 mV peak amplitude (first three spikes). In this example, the fast and slow excitatory synapse conductances and weights are equal to 1 nS and 1, respectively, creating the same strength of influence on the postsynaptic neuron. However, the combination of the neuronal and astrocytic inputs triggers a sudden increase in *v* up to −46.35 mV. Then, *v* decays to 0 while fluctuating, resulting from the fast activation of presynaptic input.

**Fig. 5:**
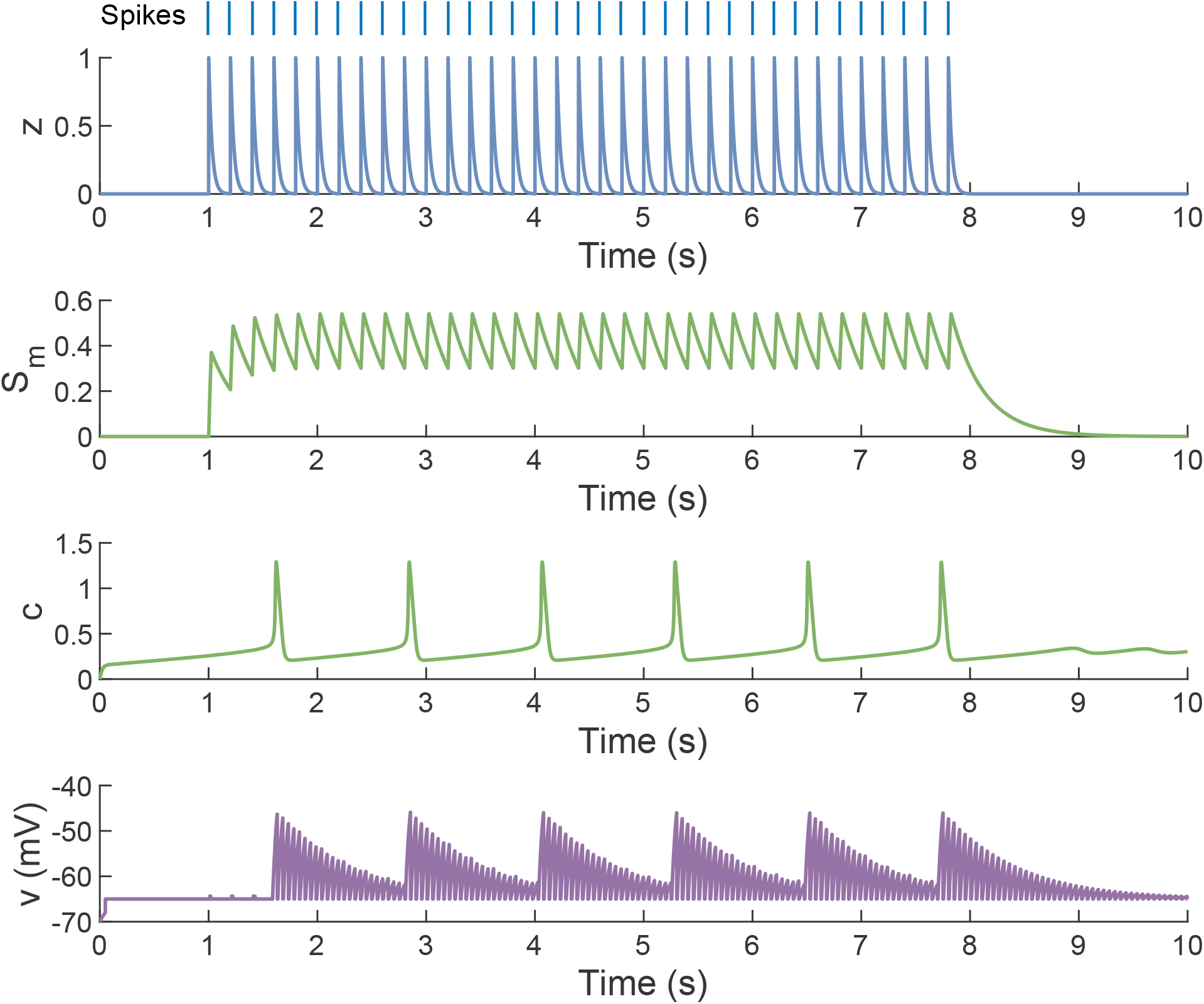
Sample tripartite synapse dynamics. (Parameters: *dt* = 1 ms, input spike rate = 5 Hz, *v*_*rest*_ = −70 mV, *θ* = −40 mV, *c*_*thresh*_ = 0.40, *g*_*e*1_ = 1 nS, synaptic delay = 9.62 ms, *β* = 0.006, *τ*_*c*_ = 50 ms, *g*_*e*2_ = 1 nS.)

In the SNAN, astrocytes receive hundreds of inputs; therefore, the summation of slow synaptic weights must generate a conductance, *g*_*exc*2_, low enough to not overexcite the postsynaptic neuron. In addition, the neuron-astrocyte coupling control factor *β* must be high enough to trigger astrocytic activation. We performed multiple simulations and determined the astrocytic hyperparameters in Table 3. Here, the astrocyte-neuron coupling, *w*_*exc*2_, is updated upon the arrival of astrocytic input; however, there is no *w*_*exc*2_ update (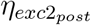is null) upon postsynaptic neuron spiking due to the significant time scales difference between the astrocyte and the neuron. If STDP is bidirectional, then the fast-spiking of the postsynaptic neuron can cancel the astrocyte-neuron coupling. High slow excitatory synaptic weights excite the First layer neurons that can cause erratic spiking patterns, suggested by the average spiking rates of SNAN2 in Figure 6a that grow with an extended number of tripartite connections. The astrocytic population rates of SNAN1 and SNAN2 (Figure 6b) have the same trend due to the similarity in *β*. For *β* = 0.001 and *β* = 0.0005 with 10% and 20% tripartite synapses, respectively, are inadequate to activate the astrocytes.

**Fig. 6:**
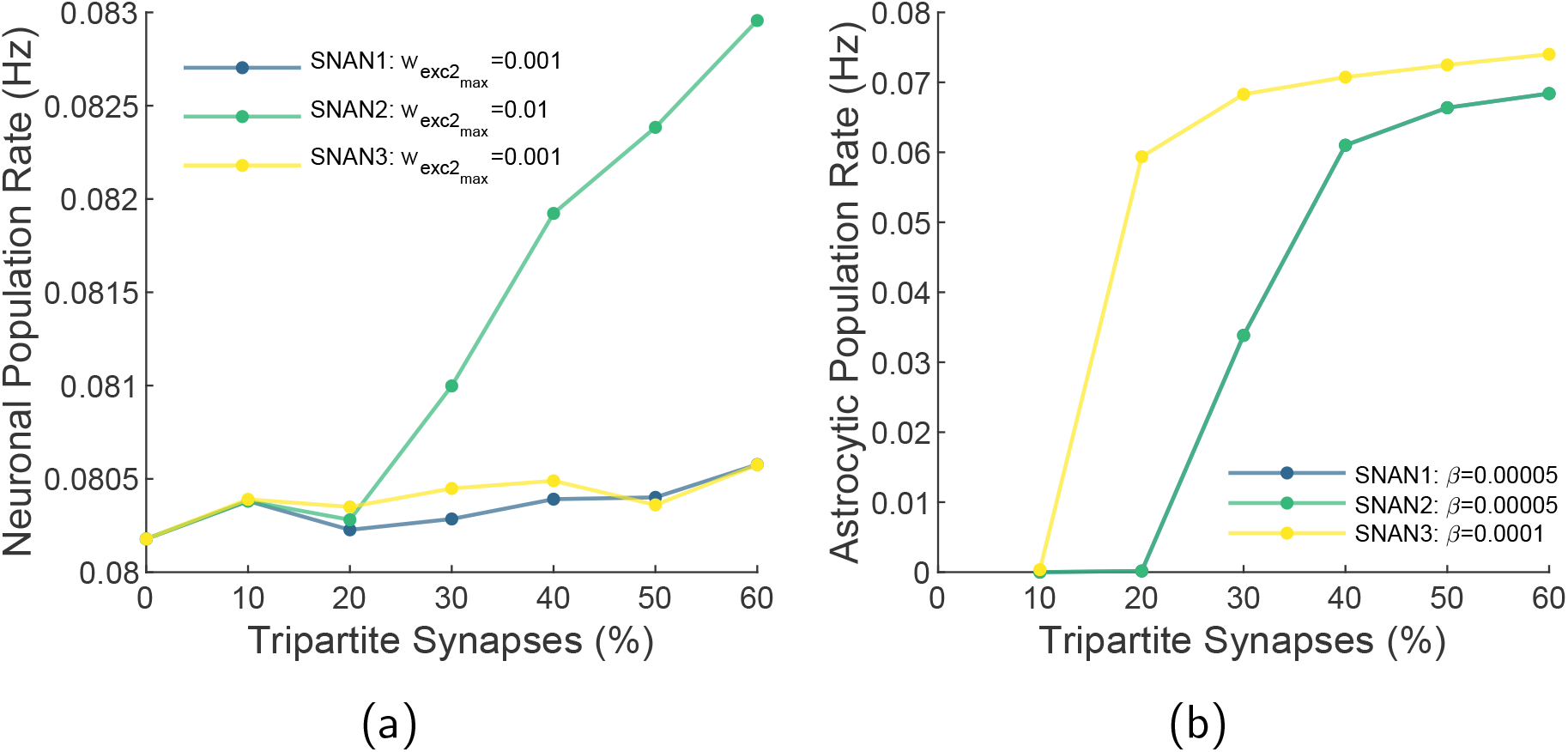
Population rates of (a) First layer neurons and (b) astrocytic layers with an increasing number of tripartite synaptic connections.

### 3.2 Training Results

We trained the networks by presenting the entire 60 000 MNIST training images for a duration spanning 25 epochs. The synaptic weight evolution in Figure 7 show the ability of the First layer neurons to learn the representative inputs. The patterns suggest which input neurons were inactive and strongly coupled with the first layer neurons. Here, the 784 excitatory-to-excitatory synaptic weights vector of a neuron is rearranged to a 28*×* 28 matrix. For example, the second neuron in the SNN learned the features of class “1”, and the ninth neuron in SNAN3 initially learned class “8” features, then became strongly coupled with class “1” inputs. The receptive fields show that neurons in the three SNAN configurations learn almost the same features, while the neurons in SNN learn differently.

**Fig. 7:**
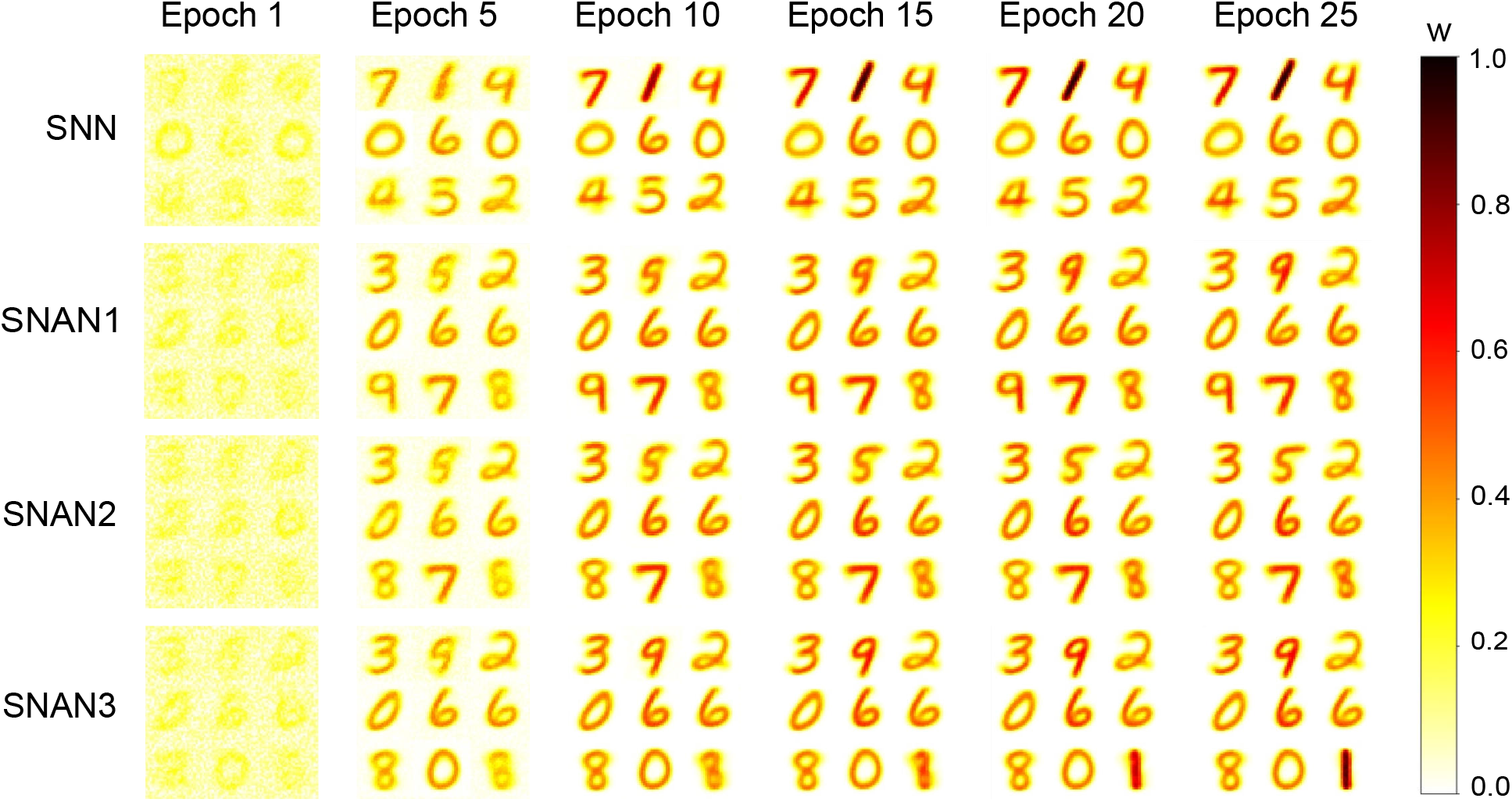
Receptive field evolutions for the nine First layer neurons of the SNN and SNANs with 50% tripartite synaptic components.

The classifier then predicts the output class based on the spiking patterns given by the SNAN per individual input. Figure 8 to summarize the change in training accuracy of each network configuration with increasing synapses covered by astrocytes. The curve with 0% tripartite synapses is the training accuracy of SNN, used as the reference in comparing the SNAN performance. The accuracy curves are the same during initial training, and the SNN gradually outperforms the SNANs. In SNAN2, it is noticeable that the rate of increase in the accuracy of SNN is maximum. However, starting from Epoch 13, the SNN performance decreases to a minimum and becomes stable at an average accuracy of 65.38%, and then all SNANs achieve higher accuracy than SNN. Figure 7 also indicates that SNANs with 50% tripartite synapses have higher performance in all network configurations, wherein the maximum accuracy is 69.94% at Epoch 13 for SNAN1, 69.76% at Epoch 25 for SNAN2, and 70.13% at Epoch 23 for SNAN3. From these results, SNAN3 with 50% tripartite synapses leads to maximum classification performance. It is also indicative that the SNAN produces spiking patterns easily categorized by the classifier unit.

**Fig. 8:**
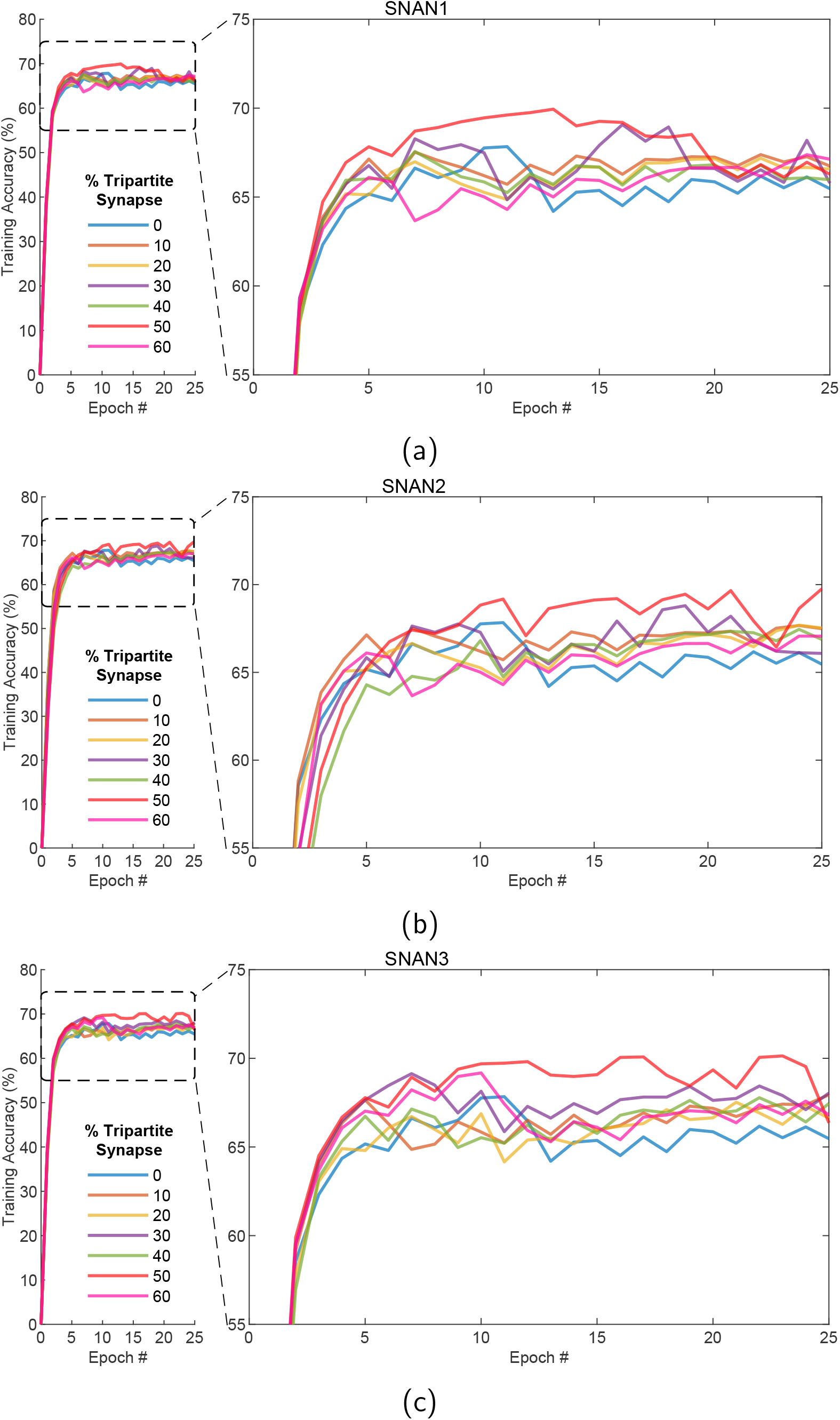
Training accuracy per network configuration (a) SNAN1, (b) SNAN2, and (c) SNAN3 with an increasing number of tripartite synapses as a function of the presented training epoch.

The validation accuracy curves, on the other hand, indicate the reverse–showing that the SNN best predicts the output class with maximum accuracy of 70.66% at Epoch 25. In SNAN1 (10%-50% tripartite synapses), the accuracy curves stabilize at an average level equal to 61.60%, almost 9% lower than SNN. In SNAN2, where the astrocyte-neuron synaptic coupling strength is a factor of 10 greater than SNAN1, the increase of tripartite connections to 50% and 60% results in a sudden decrease in validation accuracy, implying overfitting of the trained SNAN2 and that the introduction of unknown samples generates irregular spiking patterns. SNAN3 has the same coupling strength as SNAN1 but double the presynaptic-to-astrocytic coupling strength. In this case, the validation accuracy increases gradually but results in higher performance (for 20% and 30% tripartite synapses).

We chose the test network parameters from the train vs. validation accuracy graphs in Figure 9 to 11. Strikingly, the SNN network has a validation accuracy curve of 6% higher than the training accuracy, indicating under-fitting. The increase of astrocytic coverage by up to 30% diminishes the difference between the train and validation accuracy of SNAN1 (Figure 9). A further increase to 60% results in considerable divergence between the accuracy curves. The same results are noticeable in SNAN2 (Figure 10); validation accuracy increases with more astrocytic-mediated synapses but drops significantly to 10% accuracy for 60% tripartite connections. However, SNAN3 validation accuracy, with slow astrocyte-to-neuron STDP, and stronger neuron-to-astrocyte coupling (*β* = 0.001), also shows a slower convergence with the training accuracy (Figure 11). The SNAN3 with 20% and 30% tripartite connections have comparable training and validation accuracy levels in all configurations. Also, for the given SNAN3 parameters with 40%-60% astrocytic mediated synapses, the network performance improves gradually, compared to the low validation accuracy in SNAN2 (Figure 10). These results are accordant with the results of the biologically plausible model suggesting that astrocytes can either improve or impair network activities [37–39].

**Fig. 9:**
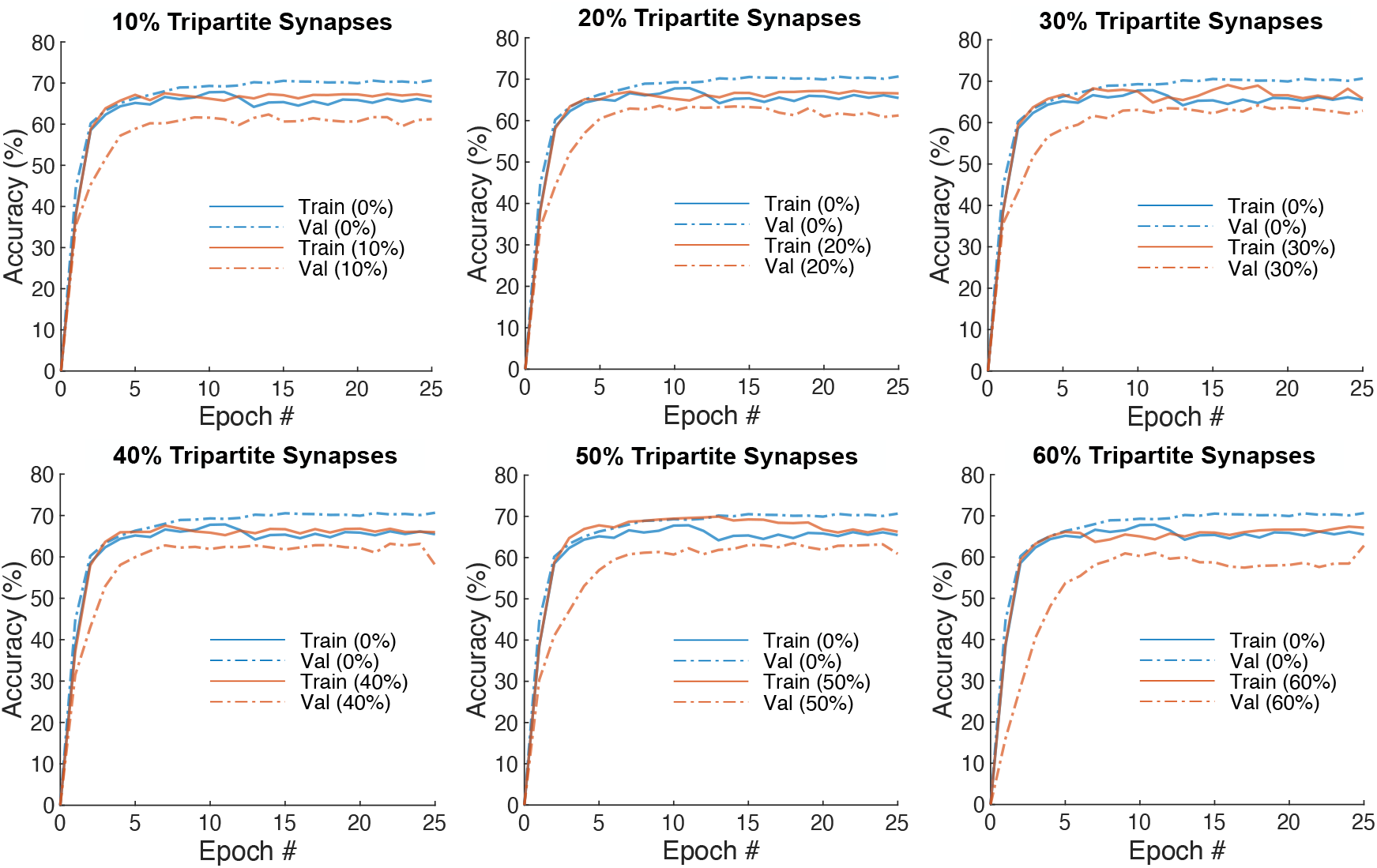
SNAN1 training (*solid lines*) and validation (*dashed lines*) accuracy for n% tripartite synapses. (Parameters: 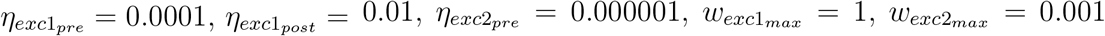, *β* =0.00005.)

**Fig. 10:**
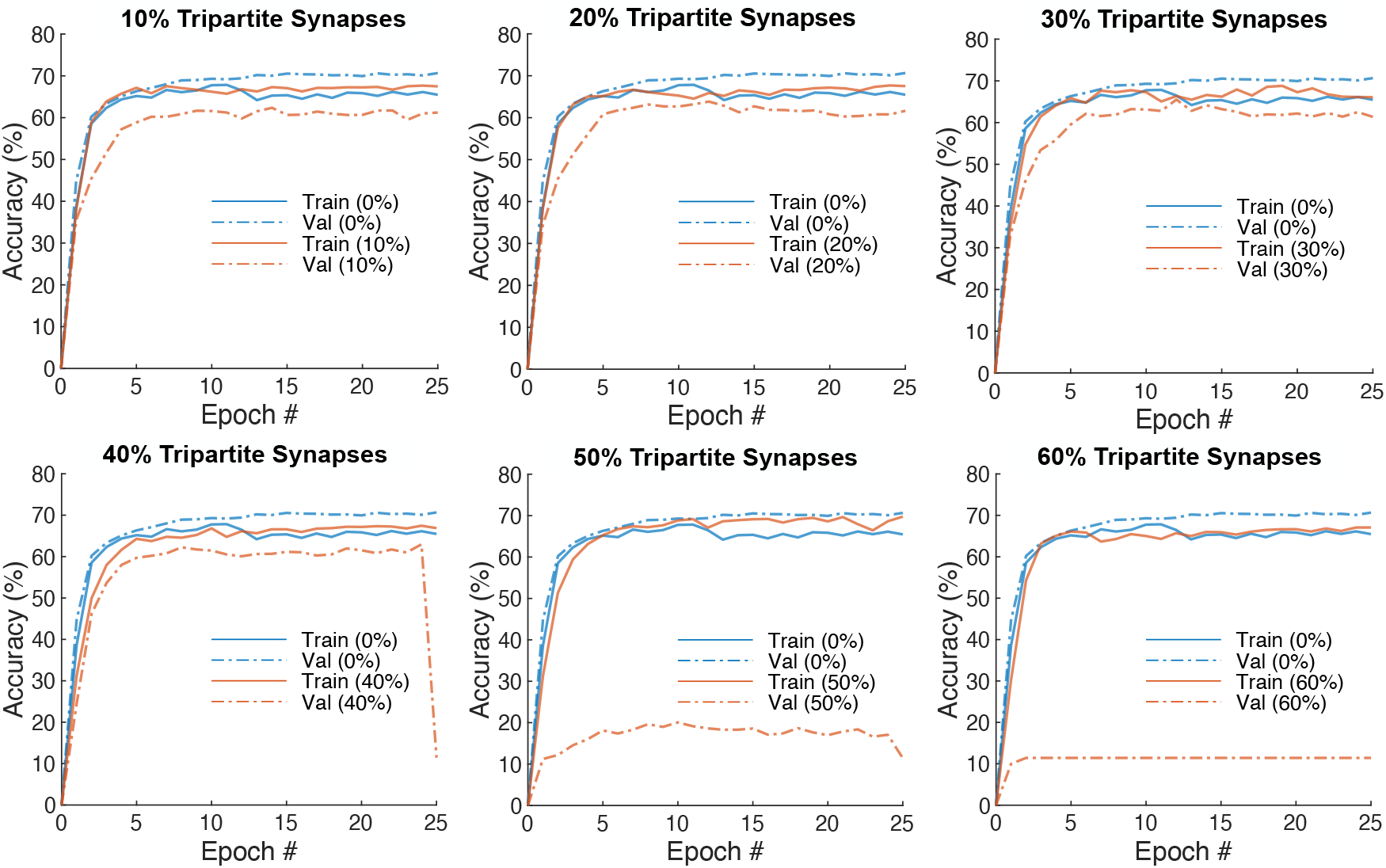
SNAN2 training (*solid lines*) and validation (*dashed lines*) accuracy for n% tripartite synapses. (Parameters: 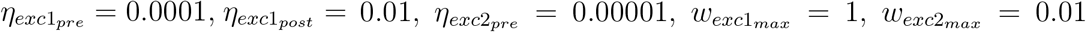, and *β* =0.0005.)

**Fig. 11:**
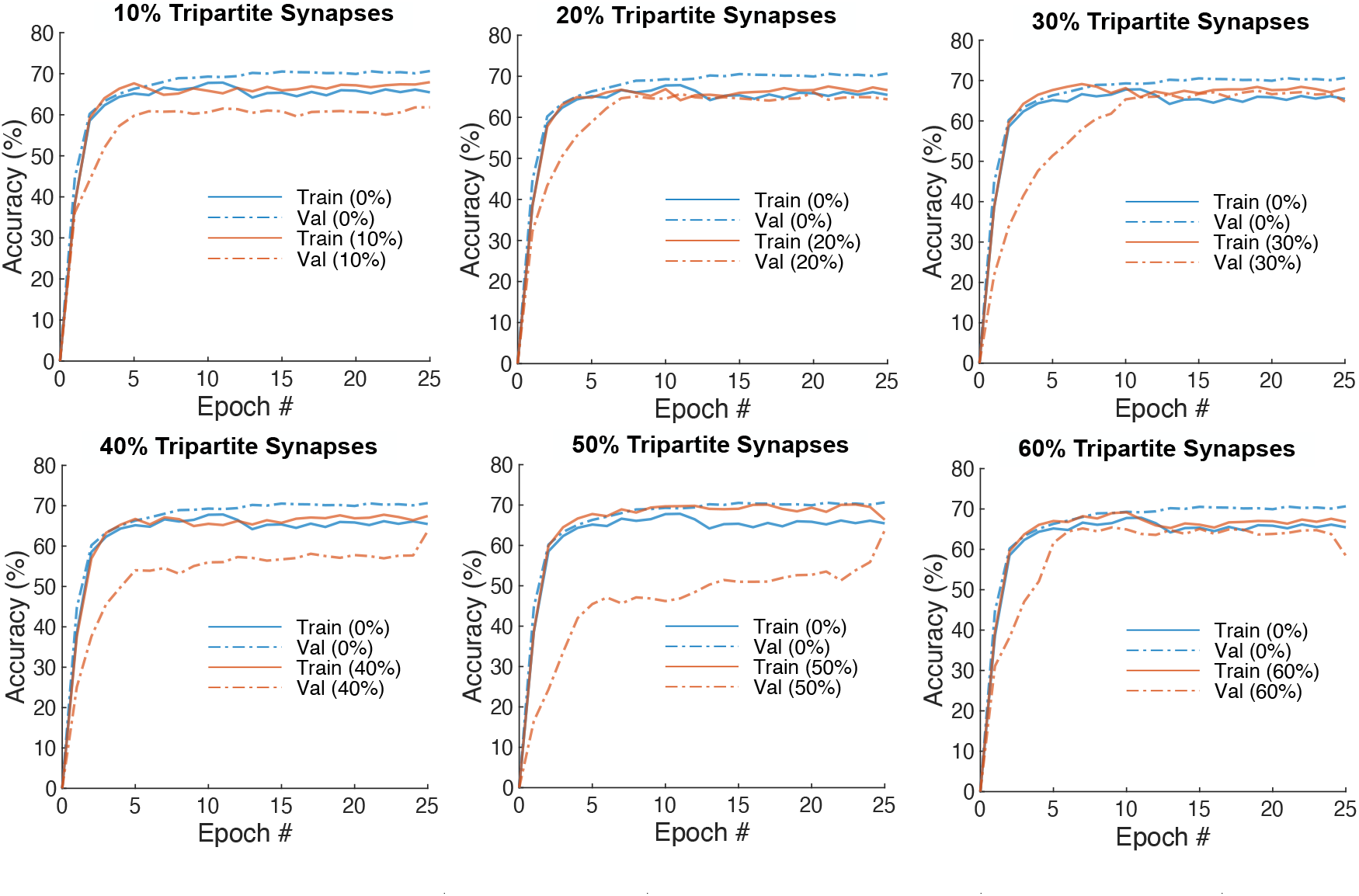
SNAN3 training (*solid lines*) and validation (*dashed lines*) accuracy for n% tripartite synapses. (Parameters: 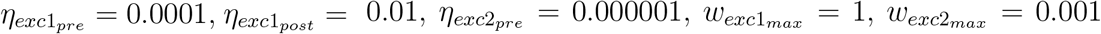, and *β* = 0.001.)

The network parameters giving the optimal model complexity (balance between variance and bias) are the learned parameters after the epochs specified in Table 4. The SNN is optimal after ten times input presentations.

**Tab. 4:** Epochs of the network parameters with optimal model complexity.

| % Tripartite Synapse | SNAN1 |  | SNAN2 |  | SNAN3 |  |
| --- | --- | --- | --- | --- | --- | --- |
|  | Train Accuracy (%) | Epoch # | Train Accuracy (%) | Epoch # | Train Accuracy (%) | Epoch # |
| 0 | 67.76 | 10 | 67.76 | 10 | 67.76 | 10 |
| 10 | 67.30 | 14 | 67.30 | 14 | 67.94 | 25 |
| 20 | 67.12 | 19 | 65.90 | 12 | 66.65 | 20 |
| 30 | 68.94 | 18 | 66.34 | 12 | 68.41 | 19 |
| 40 | 67.56 | 7 | 67.45 | 24 | 67.45 | 25 |
| 50 | 69.26 | 15 | 68.83 | 10 | 66.36 | 25 |
| 60 | 67.13 | 25 | 67.07 | 24 | 68.97 | 9 |

The SNN, in this case, displays underfitting (having a higher validation than training accuracy). SNANs achieve higher network performance and, in some instances, require less training. For example, it only takes seven iterations for SNAN1 with 40% tripartite connections to achieve an accuracy of 0.20% less than SNN, and nine iterations for SNAN3 with 60% tripartite connections to obtain 1.21% higher than SNN. In addition, for the same number of epochs, SNAN2 with 50% tripartite connections achieves 1.07% higher than SNN. The results in Table 4 suggest that astrocytes support neuronal memory formation and promote faster learning.

### 3.3 Test Results

We tested the network using the 5000 test images from the MNIST dataset. The test simulations take an average of 0.41 s, 0.69 s, 0.88 s, 1.02 s, 1.33 s, 1.67 s, and 1.59 s for the networks with 0%, 10%, 20%, 30%, 40%, 50%, and 60% tripartite synaptic connections, respectively, to predict the class of a single test image. Then, Table 5 shows the resulting test accuracy per network configuration. The proposed SNN gives an accuracy of 82.46%, 0.44% less than the accuracy reported by Diehl and Cook [30] for their network with 100 First layer neurons. Of the neuron-astrocyte networks, the parameters of SNAN3 give the best network performance, with maximum accuracy of 75.28% for 30% tripartite connections. Indeed, the neural network displays higher prediction accuracy than the neuron-astrocyte networks. However, the neuron network also displays high bias and underfitting, given that the test accuracy is 14.17% greater than the training accuracy. It suggests the SNN network is not yet fully optimized and requires extended training for more than 25 epochs. Therefore, the astrocytes improve network performance, especially in the SNAN3, where there is an optimum variance-bias tradeoff.

**Tab. 5:** Test accuracy.

| % Tripartite<br>Synapse | SNAN1 | SNAN2 | SNAN3 |
| --- | --- | --- | --- |
| 0 | 82.46 | 82.46 | 82.46 |
| 10 | 65.46 | 65.46 | 66.80 |
| 20 | 67.54 | 66.44 | 71.96 |
| 30 | 68.94 | 69.98 | 75.28 |
| 40 | 67.26 | 67.32 | 70.26 |
| 50 | 68.56 | 21.66 | 72.44 |
| 60 | 71.20 | 11.28 | 71.96 |

The diagonals of the confusion matrices in Figure 12 shows the number of correctly predicted input class. Even though the SNN correctly predicted class “0” more times than the SNAN, the SNAN still generates higher precision (92%) for the specified class, given that there are more instances that the SNN confuses the remaining input classes as “0”. The SNAN precisely predicts input classes “1” (93.70%) and “3” (97.10%). Furthermore, The SNAN also displays higher recall for input classes “4” (79.90%) and “6” (95.60%), correctly predicting the input class and not confusing them with other classes. For other instances otherwise, the SNN displays higher precision and recall compared to the SNAN as expected. SNAN3 also predicts input classes “1” and “2” as “8”. This prediction can be caused by the receptive field of the neuron showing combined features of “1” and “8”, for example, the ninth neuron in the SNAN3 shown in Figure 7. SNAN3 also confuses class “9” as “4” due to their similar vertical features. The neurons with receptive fields of “9” spike at a high rate when the number “4” is presented to the network.

**Fig. 12:**
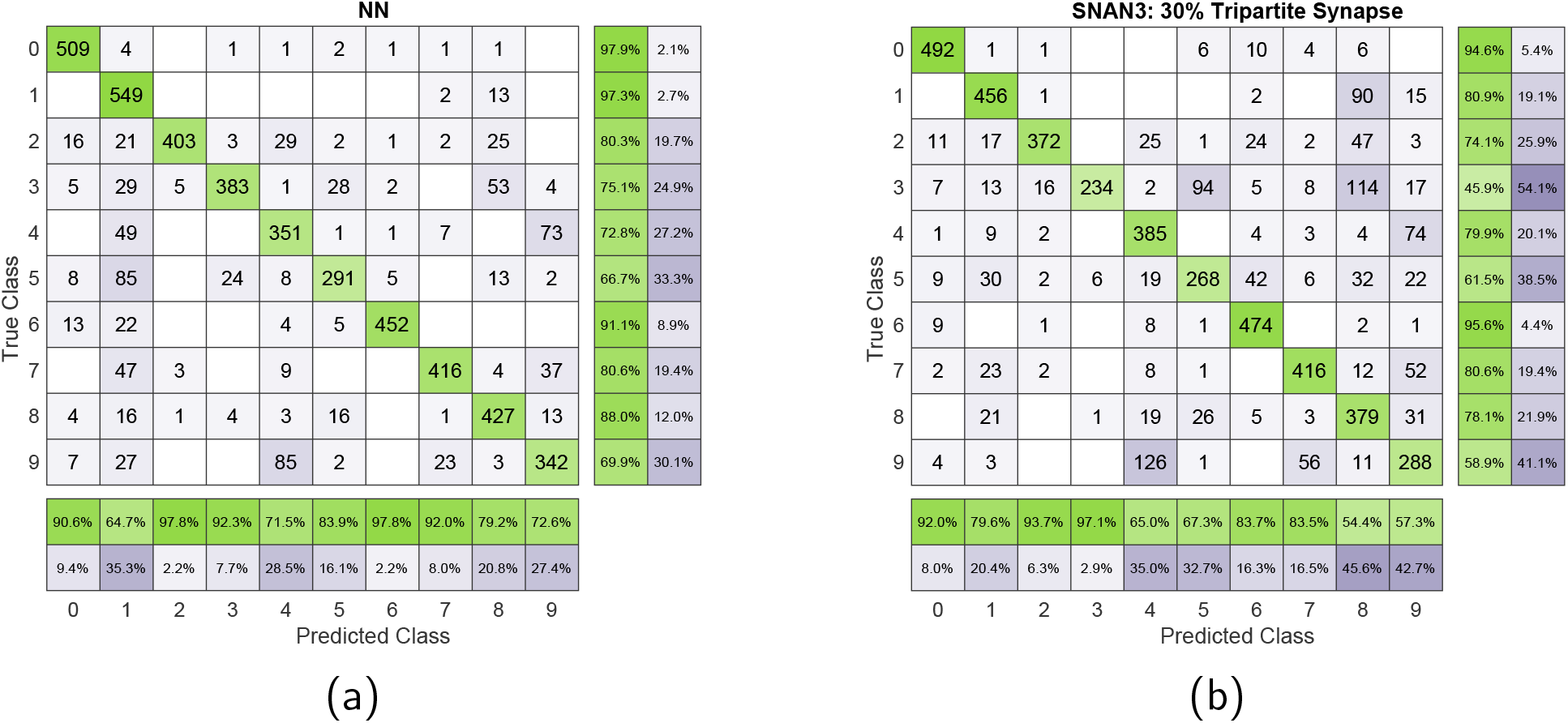
Confusion matrices for (a) SNN and (b) SNAN3 with 30% tripartite synapses. The precision and recall are displayed, respectively, below and on the right of the confusion matrix.

## 4 Discussion

We proposed a novel and simplified spiking neuron-astrocyte network for unsupervised learning using STDP, combined with k-NN classifier for image recognition. The input layer consists of Poisson spiking neurons translating the input image pixel intensity into spike events. The dense synaptic connections transmit the input activities to the 100 postsynaptic neurons. The postsynaptic neurons then learn the input features by changing their synaptic weights following the STDP algorithm. In these networks, astrocytes have three main functions: (1) create a new network layer for integrating and translating presynaptic inputs, (2) act as additional synaptic connections simplifying network architecture, and (3) modulate synaptic plasticity for faster learning. We have achieved a maximum of 75.28 % of classification performance for SNANs, ensuring an optimal balance between variance and bias. We showed that astrocytes improve network performance (depending on their topology within the network) and facilitate stable neuronal spiking.

### 4.1 Towards the Development of SNANs for Deep Learning

Furthermore, we presented a hybrid network of spiking networks and machine learning, derived from the baseline SNN by Diehl and Cook [30], where the SNAN generates a spiking pattern specific to the input class and the *k*-NN classifier predicts the output class. Diehl and Cook [30] reported a classification performance of 82.90% for 100 first layer neurons. With our proposed baseline SNN, we acquired a relative value equal to 82.46%. Our analysis suggests that even though the baseline SNN has the highest accuracy, it displays underfitting caused by insufficient training iterations shown by the difference in training and validation accuracies (Figure 9-11). For the SNN parameters of Diehl and Cook, the maximum spiking input frequency is approximately 63 Hz, while in our proposed architecture, the maximum input frequency is about 32 Hz, because spiking frequencies above 32 Hz cause oversaturation of astrocytic Ca^2+^ dynamics, preventing astrocytic activations. Consequently, synaptic weight updates are slower. However, there was no validation set in the study of Diehl and Cook [30]. We reported a maximum test accuracy of 75.28% for the SNAN, displaying optimal variance and bias tradeoff.

Nazari et al. [26] showed a 77.35% accuracy in digit classification using a cortical neuron-astrocyte network that was trained on the MNIST dataset; this was in contrast to our proposed network. Their extensive architecture had a significant number of fully connected astrocytes (1,501,674) in addition to leaky integrate-and-fire neurons (5,000), which resulted in a marginally superior performance of 2.07% over our network. However, our proposed network featured a simplified configuration of only 100 astrocytes (784 + 100 IF neurons) and showed a comparable results while benefiting from a more simplified design. On a separate work by Nazari and Faez [26], the incorporation of 2,500 astrocytes forming tripartite synapses with pyramidal neurons (2,500) and interneurons (2,500) improved the network accuracy. These astrocytes played a crucial role in mitigating errors caused by neurons being in an “always firing” state [26]. Determining the optimal count of astrocytic and tripartite synaptic connections is an important factor in designing the SNAN. Our study shows that augmenting the number of astrocytic connections does not necessarily lead to a corresponding improvement in network performance, as shown in the accuracy curves of networks with 50% and 60% tripartite connections (Figure 9-11). Although astrocytes play a critical role in preventing neuronal overexcitation, it is important to note that they can also inadvertently inhibit neuronal excitation, potentially resulting in slower learning.

Rastogi et al. [27] recently demonstrated SNAN designs using the MNIST dataset, where astrocytes modulate the presynaptic neuron release probability during faults when synaptic learning is stuck at zero. In this case, the astrocytes indirectly modulate synaptic activities. In our proposed network, astrocytes act as excitatory inputs to the postsynaptic neurons with STDP ability. To our knowledge, the proposed neuron-astrocyte network is one of the first attempts to identify the potential of astrocytes in image classification using the standard MNIST dataset. Therefore, this study can serve as a baseline for evaluating future studies on astrocyte implementation in SNNs.

### 4.2 Astrocytes Improve Network Performance

Our results suggest that astrocytes influence and improve three attributes of the spiking network: (1) faster learning, (2) variance and bias tradeoff, and (3) simplified network architecture. During training, the SNANs displayed higher accuracy than SNN (Figure 8), indicative of the ability of the network to stabilize its spiking activity quickly. Moreover, the SNANs achieved peak accuracy with fewer input set presentations (Table 4). These activities support the idea that astrocyte-mediated neuronal spiking aids in faster learning. For traditional SNN and artificial networks, increased neuronal connectivity equates to fast and precise learning. Therefore, the astrocytes and the corresponding tripartite synapse promote efficient learning by providing additional synaptic connections in parallel with the neuron-to-neuron couplings, thus strengthening the communication strength between neuronal layers. Also, as the neuron spikes more frequently, the spiking threshold increases, thus requiring more input eventually. Astrocytes, therefore, provide additional input for equalizing neuronal firing rate.

Second, networks with astrocytes balanced variance and bias, as shown in comparing the training and validation accuracies (Figure 9-11). The SNAN3 with 30% tripartite synaptic connections exhibits optimal variance and bias tradeoff, with higher training accuracy than SNN and comparable validation accuracy. These metrics ensure that the spiking patterns from the validation set matched the spiking patterns produced during training. The SNN displays underfitting, exhibited by a higher validation accuracy than training accuracy, conveying that the trained network has difficulty generalizing new data. Defining the SNAN architecture is a highly iterative process due to the heterogeneity of the astrocytes combined with the stochastic spiking activity. Therefore, the number of astrocyte-to-neuron synapses must be sufficient to influence the postsynaptic neuron spiking and low enough to avoid neuronal overexcitation resulting in erratic neuronal spiking patterns. The disruption of astrocyte function results in a disturbance in the interaction between neurons and astrocytes. Consequently, this imbalance affects synaptic transmission and leads to an elevated level of excitation within the neuronal population [25]. These effects are noticeable in the networks with more than 30% tripartite connections (Figure 10), whose validation accuracy suddenly drop to minimum levels.

Lastly, we have developed a SNAN where astrocytes are point processes with simpler models while retaining the essential astrocytic dynamics. Interestingly, astrocytes can simplify the network architecture. An SNN achieves faster learning with increasing synaptic connections, which is maximum when neurons have dense synaptic connections. If the dense connection is insufficient, another solution is increasing the number of postsynaptic neurons. Saunders et al. [40] presented a network with 100 and 400 postsynaptic neurons and reported that the training accuracy increases from 67% to 70%, respectively. Therefore, for example, a network with 400 neurons creates 316 600 synaptic connections. The proposed SNAN3 with 30% tripartite achieved 75.28% accuracy with only 78 400 synaptic connections plus 47 040 (neuron-to-astrocyte and astrocyte-to-neuron synaptic connections) with only 200 integrating components (100 neurons and 100 astrocytes).

### 4.3 Neuromorphic Neuron-Astrocyte Networks

Indeed, the SNAN architecture exhibited a lower accuracy when compared to the previous SNN models. One of the main challenges we faced during network design is the simulation costs, where simulations are restricted to series computation rather than in parallel. The additional astrocytic components plus the neuron-astrocyte couplings require more extended simulation and computationally heavy programs. Instead of using the Li-Rinzel model, we opted for the Postnov model and simplified the corresponding code using the Brian2 simulator with high-performance computing resources. Researchers can also design and implement SNAN using neuromorphic systems or hardware to solve the series computation restrictions. Two projects aim to implement astrocytes in neuromorphic chips. One project, known as BioRC, is conducted by the University of Southern California [41]. The other project is a collaborative effort between the University of Tehran and the University of Kermanshah in Iran implemented on Field Programmable Gate Arrays (FP-GAs) [42]. Additionally, the RNASA-IMEDIR group from the University of A Coruña has developed an Artificial Neuron-Glia Network (ANGN), which combines artificial neurons and artificial astrocytes as different types of processing elements [13]. This study is one of the initial attempts to incorporate astrocytes in the context of third-generation neural networks. Due to computational constraints, each configuration of the proposed neuron-astrocyte network was simulated for only 25 epochs. By advancing to hardware implementation, we will be able to extend the training process and potentially address the issue of underfitting. Therefore, we can conduct a comprehensive performance comparison using various data sets.

In summary, we demonstrate the viability of integrating astrocytes into artificial networks in conjunction with neurons, albeit with slower dynamics. In this study, we developed an astrocyte-modulated spiking neuron network optimized for image recognition. Mimicking their counterparts in biological systems, astrocytes can enhance the performance of spiking networks. However, determining optimal network performance comes with a caveat: it is crucial to find the right balance in the number of tripartite synaptic connections, which is critical for stabilizing network activity and preventing neuronal overexcitation while avoiding the risk of underexcitation. In this study, we demonstrate the potential of astrocytes to refine neural network dynamics and functionality.

## Acknowledgement

This work was partially funded by the European Regional Development Fund (ERDF) and by “Junta de Extremadura” (Ref. IB20040), and by the Ministerio de Ciencia e Innovación through grant PID2019-110315RB-I00 funded by MCIN/AEI/10.13039/501100011033 (APRISA).

Calculations were also performed using HPC resources from the Direction du Numérique - Centre de Calcul de l’université de Bourgogne (DNUM CCUB).

